# Astrocytes control cocaine-induced synaptic plasticity and reward through the matricellular protein hevin

**DOI:** 10.1101/2023.03.19.533284

**Authors:** Raphaële Mongrédien, Augusto Anesio, Gustavo J. D. Fernandes, Andrew L. Eagle, Steeve Maldera, Cuong Pham, Adèle Vilette, Paula Bianchi, Clara Franco, Franck Louis, Carole Gruszczynski, Catalina Betancur, Amaia M. Erdozain, Alfred J. Robison, Antony A. Boucard, Dongdong Li, Fabio C. Cruz, Sophie Gautron, Nicolas Heck, Vincent Vialou

## Abstract

Drug addiction involves profound modifications of neuronal plasticity in the nucleus accumbens, which may engage various cell types. Here, we report prominent effects of cocaine on calcium signals in astrocytes characterized by *in vivo* fiber photometry. Astrocyte calcium signals in the nucleus accumbens are sufficient and necessary for the acquisition of cocaine seeking behavior. We identify the astrocyte-secreted matricellular protein hevin as an effector of the action of cocaine and calcium signals on reward and neuronal plasticity.

## INTRODUCTION

Astrocytes in the nucleus accumbens (NAc) play a dynamic role in regulating glutamatergic neurotransmission during psychogenic drug experience, promoting drug seeking and relapse after drug withdrawal^1^. Enhanced glutamatergic AMPA receptor neurotransmission on NAc medium spiny neurons (MSN) synapses, and reduced astrocytic glutamate transporter GLT1 expression, causing glutamate spillover^2,3^, have been shown to promote drug seeking and propensity to relapse^1,4^. In addition to these mechanisms, other adaptive changes may be instrumental in the reprogramming of brain circuits that underlie drug seeking behavior^5,6^, notably the secretion of astrocytic factors^7^. Hevin is a matricellular protein of the SPARC (secreted protein acidic and rich in cysteine) family, predominantly expressed in perisynaptic glia processes and the excitatory post-synaptic density^8,9^ in the adult brain. Hevin exhibits synaptogenic properties that can modulate excitatory synaptic responses in neuronal cultures^8,10,11^ and modulates the plasticity linked to motivation and behavioral adaptation to adverse events^12^. Here, we define the role of astrocyte calcium (Ca^2+^) signals *in vivo* and hevin in mediating the rewarding effects of cocaine.

## RESULTS

Astrocyte Ca^2+^ signals occurring *in vivo* in the NAc in response to reward cues have not been finely characterized. We monitored astrocyte Ca^2+^ signals in the NAc of freely behaving mice in response to cocaine using fiber photometry^13^, which enables temporal recording of cell population activity (Fig 1a). Ca^2+^ signals were measured using the genetically encoded Ca^2+^ indicator GCaMP6f selectively expressed in NAc astrocytes by adeno-associated viruses (AAV2.5) with an astrocyte-specific GfaABC1D (GFAP) promoter (Fig 1b). Spontaneous astrocyte Ca^2+^ signals were of relatively high amplitude and low frequency (Fig. 1c), suggesting phasic synchronization. Compared to saline, injection of cocaine (10 mg/kg, i.p.) significantly reduced peak amplitude while increasing the frequency of Ca^2+^ signal oscillations (Fig. 1d). This shows that cocaine modulates astrocyte Ca^2+^ signals in the NAc, possibly by decreasing their synchronization or amplitude, or both.

**Fig. 1.**
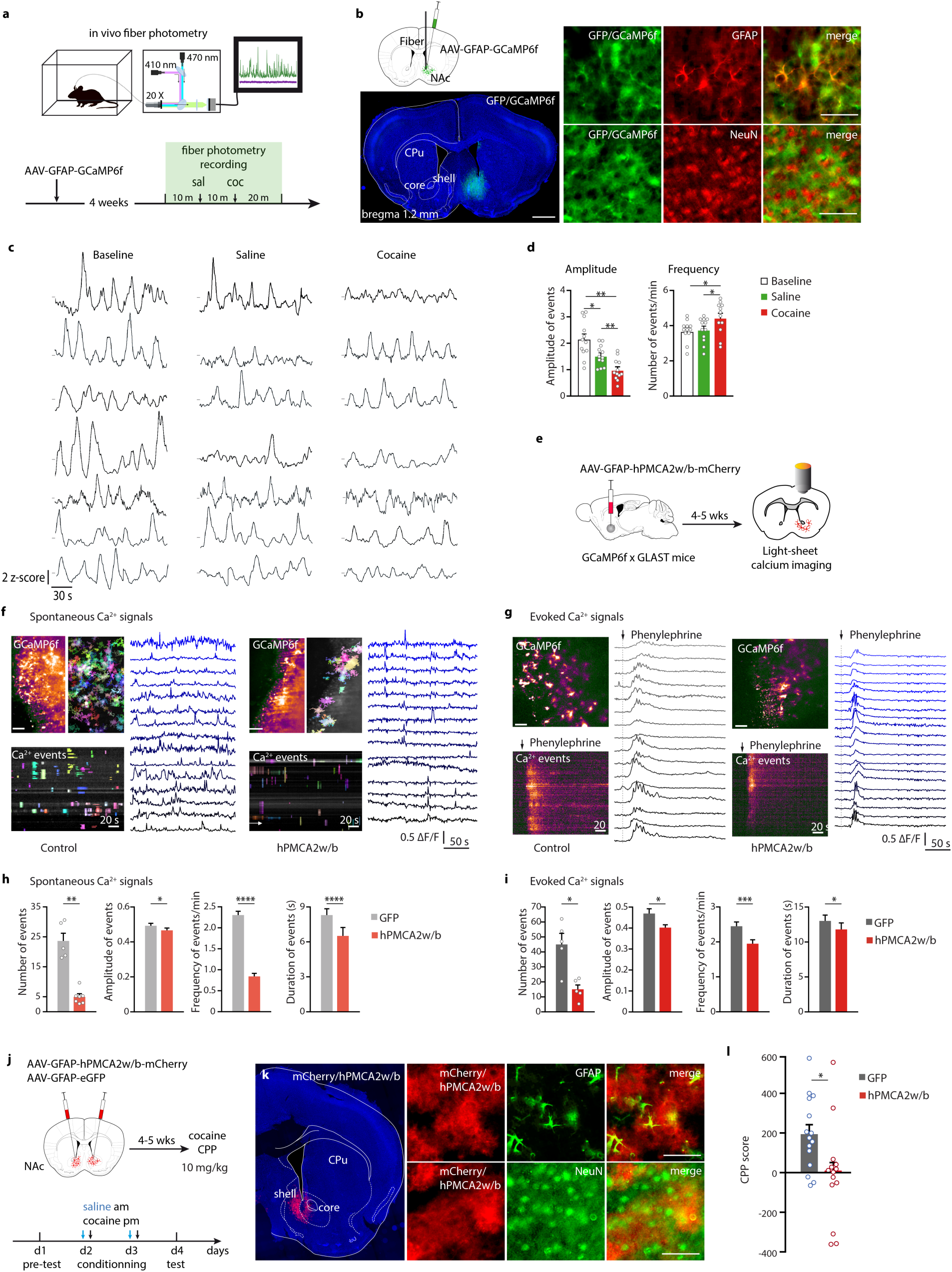
Astrocyte Ca^2+^ signals are required for cocaine CPP. **a**, Set up and timeline of the photometry experiments. **b**, Schematic representation of AAV-GFAP-GCaMP6f injection and optical fiber placement in the NAc and representative image of GCaMP6f expression at injection site. Scale bar, 1 mm (left). Representative image of GCaMP6f expression in GFAP-expressing cells (top right); minimal expression of GCaMP6f in NeuN-expressing neurons (bottom right). Scale bar, 50 μm. **c**, Example of Ca^2+^ signal z-score of deconvoluted activity traces after habituation, saline injection and cocaine injection. **d**, Average Ca^2+^ signal amplitude (left) and frequency (right) during habituation, and after saline or cocaine injection. *n* = 12. One-way repeated measures ANOVA followed by Tukey’s post hoc tests. **e**, Representation of the light-sheet imaging experiment. **f**, Spontaneous astrocyte Ca^2+^ activity in control (left) or hPMCA2w/b (right) conditions. Example of the field of view depicting temporal projection of fluorescence activity and color-coded AQuA-detected Ca^2+^ events (top), and their temporal kymograph (bottom). **g**, Evoked astrocyte Ca^2+^ activity detected by light-sheet imaging in control (left) or hPMCA2w/b (right) conditions. Example of the field of view depicting temporal projection of fluorescence activity (top) and temporal kymograph of GCaMP6f-evoked responses (bottom). Scale bar, 50 μm. Traces are represented with different tonalities of blue for visualization. **h, i**, Total number of events detected, amplitude, frequency and duration of isolated spontaneous and evoked astrocyte Ca^2+^ events in control or hPMCA2w/b-injected mice (spontaneous: 373 detected events for control and 116 events for hPMCA2w/b; evoked [adrenergic GPCR stimulation]: 543 detected events for control and 375 events for hPMCA2w/b, n = 3). Two-sided Student’s *t*-test. **j**, Representation of the cocaine CPP experiment after bilateral injection of AAV-GFAP-hPMCA2w/b-mCherry or AAV-GFAP-eGFP control virus in the NAc. **k**, Representative image of hPMCA2w/b-mCherry expression in the NAc (left), coexpression with GFAP (top); and minimal coexpression with NeuN (bottom). Scale bar, 50 μm. **l**, Cocaine CPP score after bilateral injection of AAV-GFAP-hPMCA2w/b or AAV-GFAP-eGFP control virus in NAc astrocytes. *n* = 18. Two-sided Student’s *t*-test. Data are means ± s.e.m.

To determine whether Ca^2+^ signals in NAc astrocytes are required for the rewarding properties of cocaine, these signals were decreased by expressing human plasma membrane Ca^2+^ ATPase (hPMCA2w/b) selectively in astrocytes (Fig. 1e). The effect of hPMCA2w/b on astrocyte Ca^2+^ signal dynamics was validated using light-sheet microscopy and analyzed by AQuA-based event detection^14^. Expression of the Ca^2+^ pump significantly inhibited spontaneous and evoked Ca^2+^ signals in NAc astrocytes, as evidenced by a reduction in the total number of events, and in their amplitude, frequency and duration compared with the contralateral non-injected side (Fig. 1f-i and Extended Data Fig. 1). The frequency of these signals was comparable to those reported using confocal imaging on slices^7,15^ but lower than that recorded *in vivo* (Fig. 1d). To test the implication of astrocyte Ca^2+^ signals in the rewarding properties of cocaine, hPMCA2w/b or control GFP were injected bilaterally into the NAc. Mice were tested using the conditioned-place preference (CPP) paradigm following administration of cocaine (10 mg/kg; Fig. 1j, k). hPMCA2w/b-injected mice showed a decrease in CPP compared to GFP-injected control mice (Fig. 1l). These results demonstrate that Ca^2+^ signals in NAc astrocytes contribute to the rewarding properties of cocaine.

Matricellular proteins secreted by astrocytes are essential for the formation and stabilization of synapses through cell-cell interactions^16^. The matricellular protein hevin (also known as SPARC-like 1) was shown to promote synaptogenesis and synaptic transmission by increasing AMPA- and NMDA-receptor-mediated responses at excitatory synapses in neuronal cultures^8,10,11,16^, suggesting it could modulate neuronal plasticity and motivation in a drug-associated context. In the adult mouse brain, hevin protein expression established by immunofluorescence matched that of the corresponding mRNA^17^, with significant levels in cortical areas and septum as well as in the dorsal striatum and NAc (Fig. 2a). In the striatum and NAc, hevin was expressed exclusively in astrocytes and in sparsely distributed parvalbumin interneurons (Fig. 2b). Potential changes in hevin expression in these two brain areas following cocaine exposure were analyzed by immunoblotting. Hevin protein content was increased 12 h (Fig. 2c) and 24 h (Extended Data Fig. 2) after a single cocaine injection (10 mg/kg) in the NAc but not in the dorsal striatum. The consequences of hevin invalidation on the rewarding properties of cocaine were next analyzed using the CPP paradigm. Hevin-null (*Sparcl1*^*-/-*^) exhibited no labeling for hevin in the brain (Fig. 2d). Contrarily to wild-type mice, hevin-null mice did not develop CPP to cocaine 5 mg/kg (Fig. 2e), and developed CPP with a significant lower score with the dose of 10 mg/kg (Fig. 2f). Behavioral analysis of hevin-null mice revealed no effects on locomotor activity at basal state (Extended Data Fig. 3). Development of CPP after repeated exposure to cocaine relies on the induction in NAc MSN of the transcription factor ΔFosB, a splice product of the fosB gene^18^. Compared to wild-type mice, hevin-null mice showed decreased ΔFosB levels after five consecutive daily cocaine injections, reflecting decreased activation of MSN (Fig. 2g). To evaluate the contribution of hevin from NAc astrocytes to cocaine-induced reward, microinjection of AAV with the GFAP promoter was used to deliver microRNA (miR) sequences targeting hevin mRNA (Extended Data Fig. 4) into NAc astrocytes of wild-type mice (Fig. 2h, i)^19^. Co-labeling of hevin and the emerald green fluorescent protein (EmGFP) that is co-transcribed with miR-hevin revealed a 30% decrease in the number of hevin-positive cells in infected astrocytes (Fig. 2j), demonstrating partial hevin downregulation *in vivo*. This specific downregulation of hevin expression in NAc astrocytes of adult mice impaired significantly CPP to cocaine at two doses (Fig. 2k, l). These experiments demonstrate that astrocytic hevin in the NAc plays a major role in the rewarding properties of cocaine.

**Fig. 2.**
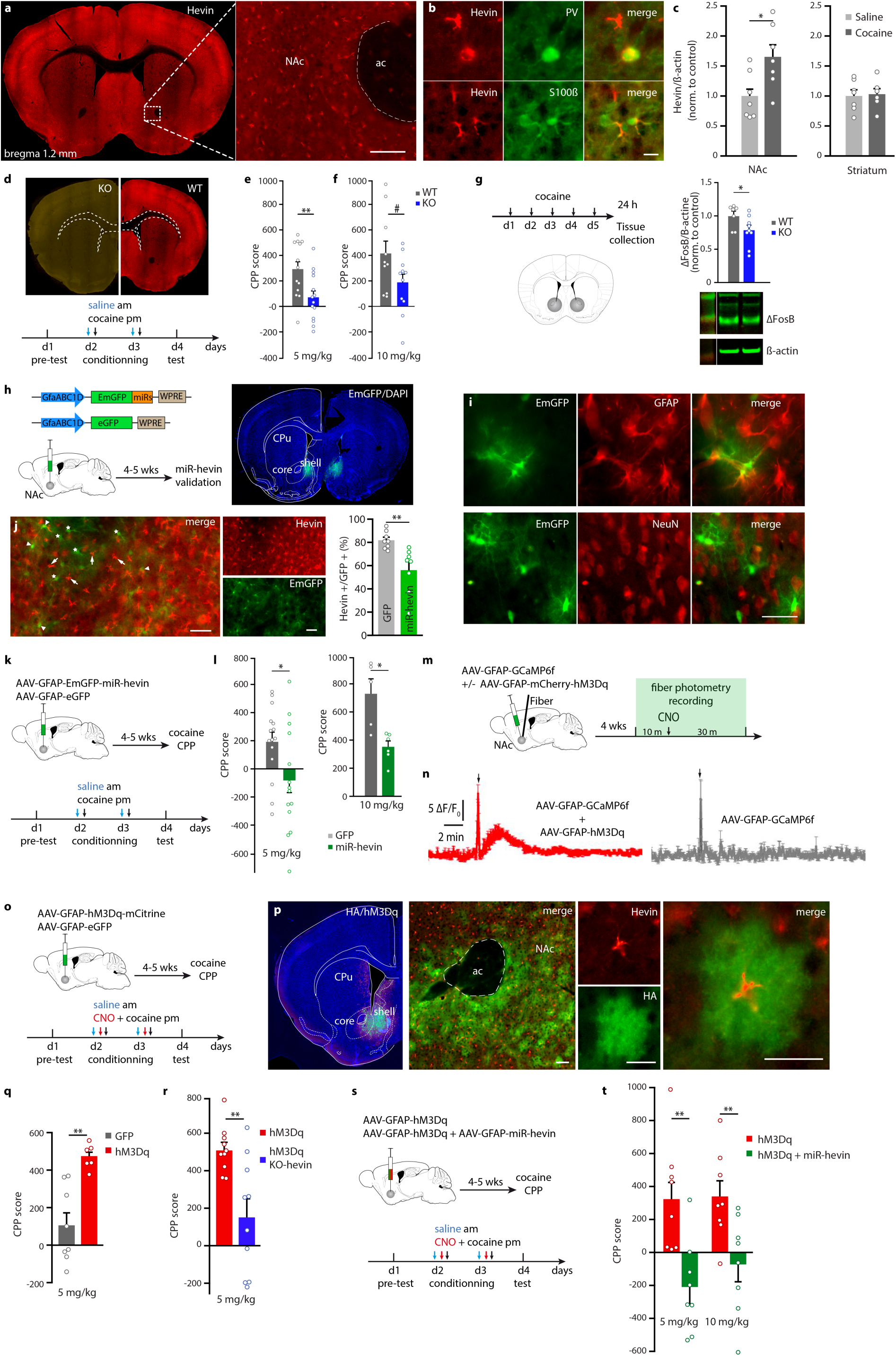
The matricellular protein hevin from NAc astrocytes contributes to the rewarding properties of cocaine. **a**, Representative image of hevin protein expression in mouse brain. Scale bar, 125 μm. **b**, Colocalization of hevin with parvalbumin and S100ß. Scale bar, 25 μm. **c**, Hevin levels in NAc and striatum after an acute injection of cocaine measured by immunoblotting. *n* = 6-7. Two-sided Student’s *t*-test. **d**, Immunohistochemistry showing lack of hevin expression in hevin-null mouse (KO) brain, and timeline of cocaine CPP. **e, f**, CPP score after low (5 mg/kg) and medium (10 mg/kg) doses of cocaine in hevin-null mice and wild-type littermates. *n* = 11-15. Two-sided Student’s *t*-test. **g**, Timeline for ΔFosB induction (left) and ΔFosB levels after repeated cocaine injection (10 mg/kg) in the NAc of wild-type and hevin-null mice (right). **h**, Schematic representation of the AAV-GFAP plasmid with EmGFP cDNA and the hevin miRNA sequences and control plasmid, and of validation procedure of *in vivo* hevin knockdown (left). Representative image of bilateral expression of EmGFP-miR-hevin restricted to the NAc (right). **i**, Representative images showing miR-hevin expression (green) colocalized with GFAP (red, top), and minimal coexpression of miR-hevin with NeuN (red, bottom). **j**, Representative images showing astrocytes expressing EmGFP-miR-hevin with (green and red, arrowheads) or without (green, stars) hevin, and non-transfected astrocytes expressing hevin (red, arrows). Scale bar, 25 μm. Quantification of hevin-positive cells in GFP-positive astrocytes (right). *n* = 6-7. **k**, Experimental procedure and timeline of cocaine CPP after bilateral injection of AAV-GFAP-miR-hevin in NAc or AAV-GFAP-eGFP control virus. **l**, CPP score after low (5 mg/kg) and medium (10 mg/kg) doses of cocaine following NAc astrocytic hevin knockdown compared to GFP-control. *n* = 14-15. Two-sided Student’s *t*-test. **m, n**, Experimental procedure and Ca^2+^ signals for validation of chemogenetic activation of NAc astrocytes (CNO, 1 mg/kg, i.p. injection; *n* = 2) using *in vivo* fiber photometry. **o**, Experimental procedure and timeline of CPP after chemogenetic activation of Ca^2+^ signals in astrocytes. **p**, Representative images of hM3Dq expression in NAc astrocytes expressing hevin. Scale bar, 25 μm. **q, r**, Cocaine CPP score after chemogenetic activation of NAc astrocytes, *n* = 6-8. Two-sided Student’s *t*-test, and in WT and hevin-null mice, *n* = 10-11. Two-sided Student’s *t*-test. **s**, Experimental procedure of cocaine CPP after bilateral co-injection of AAV-GFAP-hM3Dq and AAV-GFAP-miR-hevin or AAV-GFAP-eGFP control virus. **t**, CPP score after low (5 mg/kg) and medium (10 mg/kg) doses of cocaine associated with chemogenetic activation of NAc astrocytes after hevin knockdown compared to GFP-control. Two-way repeated measures ANOVA followed by Tukey’s post hoc tests. Data are means ± s.e.m.

To explore the importance of astrocyte Ca^2+^ signals for the rewarding properties of cocaine, human M3 muscarinic receptor (hM3Dq) designer receptor exclusively activated by designer drug (DREADD)^20^ was expressed under the control of an astrocyte promoter (AAV-GFAP-hM3Dq) into the NAc^21,22^. The exogenous ligand clozapine-N-oxide (CNO) increased the intensity of astrocyte Ca^2+^ signals detected in hM3Dq-injected mice using *in vivo* fiber photometry (Fig. 2m, n). CNO injection did not alter Ca^2+^ signals in astrocytes expressing GCaMP6f only, suggesting that it does not have off-side effects on astrocyte Ca^2+^ signals (Fig. 2n). hM3Dq-injected mice showed a notable increase in cocaine CPP score compared to GFP-injected control animals (Fig. 2o-q). This enhancement by DREADD of the rewarding effects of cocaine was dependent on hevin expression. Hevin-null mice (Fig. 2r) or mice with hevin knockdown in NAc astrocytes (Fig. 2s, t) injected with hM3Dq showed reduced rewarding effects of cocaine compared to hM3Dq-injected control mice. Altogether, these data show that enhancement of Ca^2+^ signals in astrocytes using DREADD can increase the rewarding properties of cocaine and that hevin in NAc astrocytes is required for this effect. CNO injection without cocaine pairing did not induce CPP in GFP- or hM3Dq-injected mice (Extended Data Fig. 5), demonstrating that increase of intracellular Ca^2+^ signals in NAc astrocytes did not have rewarding or aversive effects in itself.

Cocaine experience triggers long-lasting plasticity in NAc MSN that mediates in part key aspects of addiction such as motivational salience and cue-induced relapse^23^. We assessed whether hevin was implicated in the structural remodeling and reinforcement of synaptic strength of NAc MSNs after cocaine injection. Analysis of spine density of MSN in the vicinity of the bushy matrix of astrocytes that expressed EmGFP-miR-hevin or GFP-control was carried out using confocal imaging after gene-gun delivery of DiI (Fig. 3a, b). Hevin knockdown did not affect spine density at basal state (Fig. 3c). Twenty-four hour after a two-day cocaine CPP procedure, MSN spine density was significantly reduced in hevin knockdown mice (Fig. 3d) compared to GFP-injected controls. Analysis of spine length and spine head size revealed no difference between GFP-injected controls and hevin knockdown mice at basal state (Extended Data Fig. 6) or after cocaine (Fig. 3e).

**Fig. 3.**
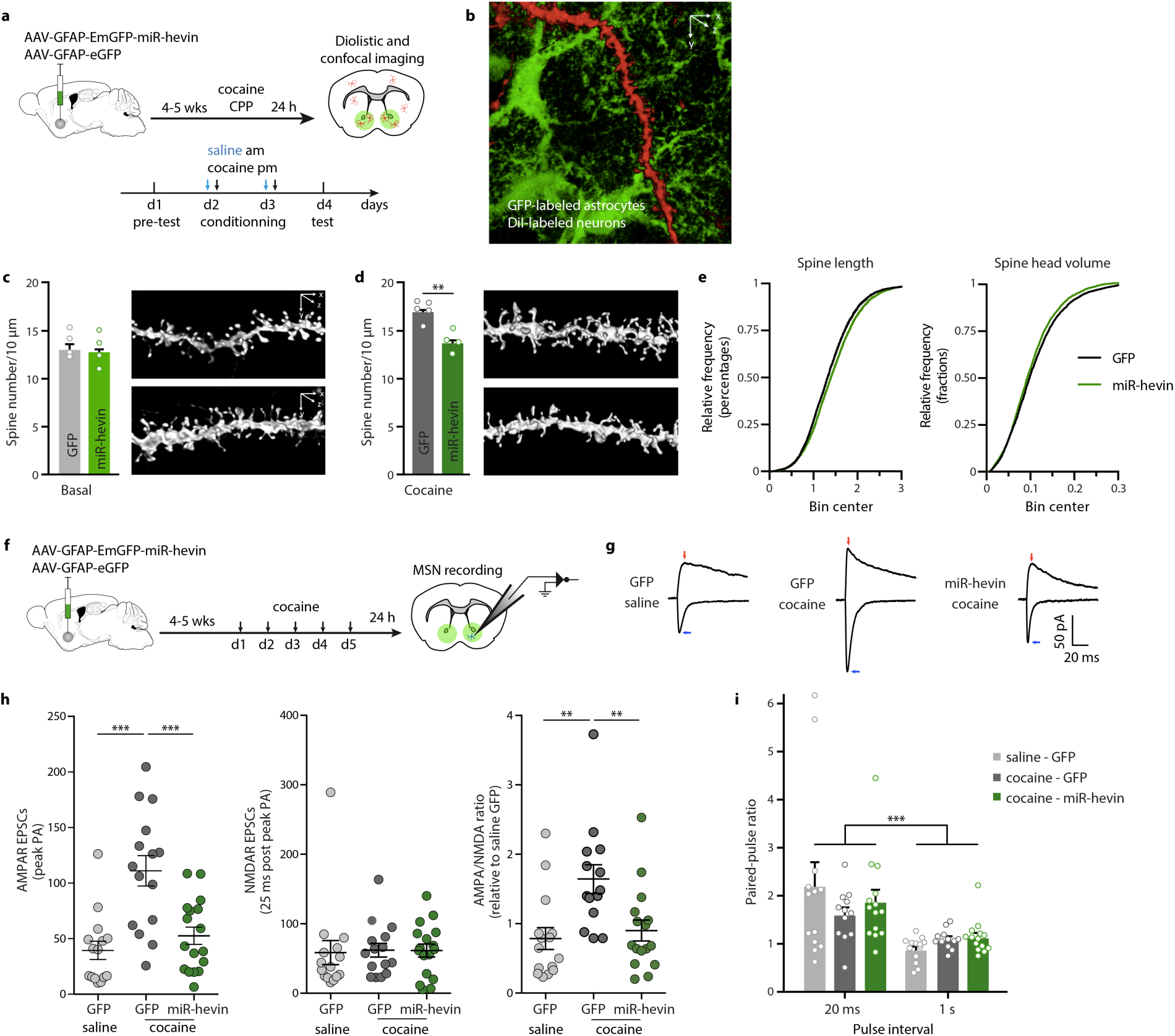
The matricellular protein hevin in NAc astrocytes contributes to cocaine-induced neuronal plasticity. **a**, Experimental procedure for confocal imaging of MSN dendritic spines after hevin knockdown in NAc astrocytes. **b**, Representative image of a DiI-filled dendrite adjacent to GFP-expressing astrocytes. **c**, Spine density in basal conditions and representative deconvoluted images of MSN dendrites after hevin knockdown compared to GFP control. *n* = 4-5. Two-sided Student’s *t*-test. **d**, MSN spine density after cocaine CPP and representative deconvoluted images of MSN dendrites after hevin knockdown compared to GFP control. *n* = 4. Two-sided Student’s *t*-test. **e**, Spine length and spine head diameter after cocaine CPP following hevin knockdown compared to GFP control *n* = 4. **f**, Experimental procedure for whole-cell patch-clamp recording of MSN after hevin downregulation in NAc astrocytes. **g**, Representative EPSCs at −70 and +40 mV from whole-cell patch clamp recording of NAc MSNs in saline-treated GFP control mice, cocaine-treated GFP control mice, and cocaine-treated miR-hevin mice. Blue arrows indicate peak amplitudes at −70 mV for measurement of AMPA-R EPSCs. Red arrows indicate current at 25 ms post artifact for measurement of NMDA-R EPSCs. **h**, AMPA-R (left) and NMDA-R (middle) mediated EPSCs, and AMPA/NMDA ratio (right). **i**, Paired-pulse ratio in GFP control and hevin knockdown (miR-hevin) mice after saline or repeated cocaine injections. *n* = 12. One-way ANOVA followed by Holm-Sidak post hoc tests. Data are means ± s.e.m.

Strengthening of glutamatergic transmission in NAc is an important feature underlying the behavioral response after repeated exposure to cocaine^23-25^. Fast excitatory postsynaptic currents (EPSCs) onto MSNs were assessed by stimulation of glutamatergic projections at baseline and after repeated cocaine injections in coronal brain slices after astrocyte hevin knockdown and GFP-control mice (Fig. 3f). As for structural plasticity, the recordings were restricted to MSNs in the bushy matrix of astrocytes expressing EmGFP-miR-hevin or GFP-control (Fig. 3f). Repeated cocaine injection increased AMPA-receptor-mediated evoked EPSCs and not NMDA-receptor-mediated evoked EPSCs (at +40 mV), resulting in potentiated AMPA to NMDA ratios in GFP-injected control mice, indicative of increase glutamatergic synaptic strength (Fig. 3g, h). Hevin knockdown in NAc astrocytes blocked cocaine-induced AMPA-receptor-mediated EPSCs, compared to GFP-control mice injected with cocaine (Fig. 3g, h), while NMDA-receptor-mediated EPSCs were not altered by hevin astrocytes knockdown (Fig. 3g, h). The AMPAR/NMDAR ratio, which provides a measure of synaptic strength, was increased after repeated cocaine injection in GFP-control mice and this increase was hampered by hevin knockdown in NAc astrocytes (Fig. 3g, h).

To test whether astrocytic hevin knockdown could affect pre-synaptic glutamatergic release, a paired-pulse stimulation protocol was applied. Paired EPSCs evoked at −70 mV with inter-stimulus intervals of 20 ms showed increased paired pulse ratios (PPR) compared to PPR at intervals of 1 s, indicating synaptic facilitation (Fig. 3i) and likely a low probability of glutamate release onto MSNs^26^. However, neither cocaine nor astrocytic hevin knockdown affected PPR. These results suggest normal pre-synaptic glutamate release in hevin knockdown mice. In addition, hevin knockdown in NAc astrocytes produced no changes in intrinsic membrane properties compared to GFP controls, i.e., in input resistance, resting membrane potential, membrane capacitance, neuronal excitability, firing threshold, or peak and rise of action potentials (Extended Data Fig. 7). Taken together, these findings suggest that astrocyte hevin is critical *in vivo* for the reinforcement of post-synaptic strength at glutamatergic synapses after repeated cocaine exposure.

## DISCUSSION

In summary, our data reveal the implication of astrocyte activity and the synaptogenic factor hevin in cocaine-induced neuronal plasticity and behavioral adaptations. This study provides the first evidence for a role of hevin in experience-dependent plasticity in adulthood. We found that an evoked intracellular Ca^2+^ rise in astrocytes promoted the learned association between cocaine and a conditioned environment, and that Ca^2+^ signals were required for this association to occur. Fiber photometry in the NAc of freely moving mice revealed high-amplitude low-frequency spontaneous astrocyte Ca^2+^ signals, reflecting phasic synchronization of Ca^2+^ signals.

Cocaine exposure profoundly modulated the characteristics of these Ca^2+^ signals, reducing their amplitude while increasing their frequency, suggesting a redistribution of Ca^2+^ activity across the astrocyte volume. Striatal astrocytes *ex vivo* show distinct types of Ca^2+^ signals depending on the subcellular compartment, with local waves and microdomains restricted to branches and thin processes, and global waves encompassing the soma and major branches^21,27^. We speculate that cocaine attenuates spontaneous high-amplitude somatic Ca^2+^ signals while increasing local waves and microdomains exhibiting Ca^2+^-mediated events. A similar decrease in signal amplitude^28^ and increase in the number of locations showing Ca^2+^ events^7^ have been reported *ex vivo* by GCaMP6f imaging in NAc slices, after cocaine self-administration or cocaine perfusion, respectively. Ca^2+^ microdomains, which have been associated with individual excitatory synapses^29^, could be increased by glutamate and dopamine receptor activation^15,30,31^, during cocaine experience.

An increase in spine density within NAc MSNs has been positively correlated with conditioned response to cocaine^32^, and the potentiation of excitatory synapses in dopamine D1 receptor in MSN causally linked to cocaine-induced behaviors^23,24^. Astrocyte-secreted hevin seems essential *in vivo* for the structural and synaptic plasticity induced in the NAc by cocaine. A role in synaptogenesis after repeated cocaine injection was also reported previously for the astrocyte-secreted factor thrombospondin 2 and its receptor^7^. Thrombospondin 2 promotes the formation of immature silent synapses lacking AMPA-receptor in the NAc, which are implicated in incubation of cocaine craving^33^. In contrast, hevin is required *in vivo* for the formation of functional MSN synapses that become biased with high AMPA to NMDA EPSC ratios following cocaine exposure. By identifying a novel mechanism in cocaine-dependent plasticity, we provide a better understanding of the factors that influence the path to addictive behavior.

## Acknowledgements

This work was supported by funds from the Institut National de la Santé et de la Recherche Médicale (INSERM), Centre National de la Recherche Scientifique (CNRS), and Sorbonne Université, and by grants from the Brain & Behavior Research Foundation (NARSAD Young Investigator Award to VV, #17566), FP7 Marie Curie Actions Career Integration Grant (FP7-PEOPLE-2013-CIG 618807 to VV), Promouvoir l’Excellence de la Recherche à Sorbonne Université (PER-SU 2014 to VV), Agence Nationale de la Recherche (ANR JCJC 2015 ANR-15-CE16-0005, Hevinsynapse to VV), and Fundação de Amparo à Pesquisa do Estado de São Paulo (FAPESP 2019/17065-4 and 2021/12978-1 to PCB, 2018/14153-7 and 2021/07134-9 to AA, 2018/15505-4 to FCC). We thank Herb Covington for helpful discussions and comments on the manuscript; Cagla Eroglu for the hevin-null mice; Helene Sage and Gail Workman for Sc1 constructs; Mélissa Desrosiers, Camille Robert, and the AAV production facility at Institut de la Vision for viral production and purification; Stéphane Fouquet and Marie-Laure Niepon of the Imaging Facility at Institut de la Vision; Christine Mouffle and Fabrice Machulka for expert assistance in animal care and genotyping and Andy Saint-Auret for technical assistance on molecular biology.

## Author contributions

V.V. conceived the project, acquired the funding, supervised students, performed experiments and analyzed data. V.V. and S.G. wrote the paper with feedback from N.H., C.B. and D.L. A.A.B., C.B., F.C.C., N.H., D.L., A.J.R., S.G. contributed to conceptualization and the experimental design. F.C.C. contributed to funding acquisition. R.M., A.A., G.J., A.L.E, S.M., C.P., A.V., P.B., C.F., F.L., C.G., A.M.E., and V.V. performed and analyzed the experiments.

## Extended Data

**Extended Data Fig. 1.**
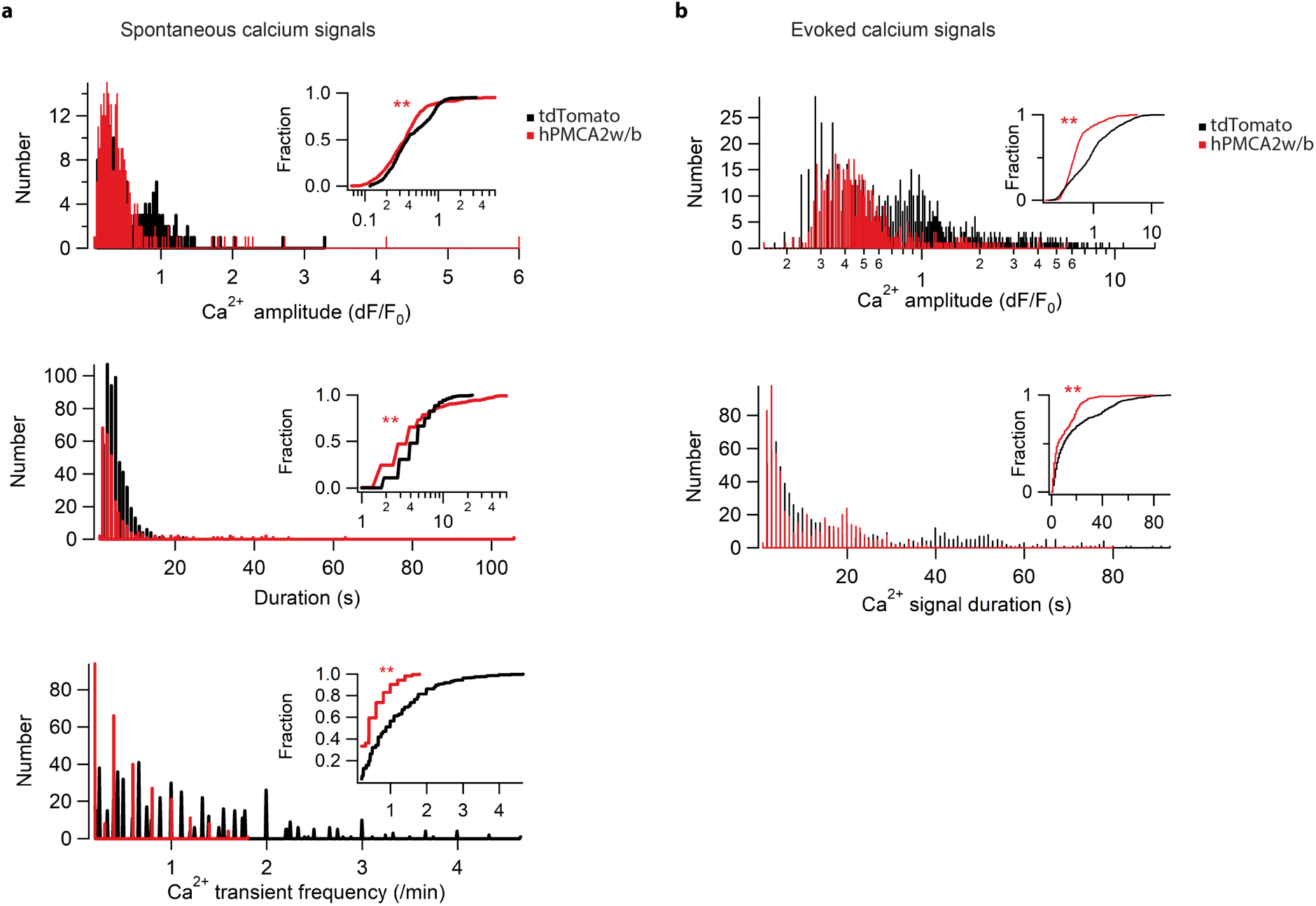
Validation of inhibitory effect of hPMCA in mice brain slices. **a**, Spontaneous astrocyte Ca^2+^ activity in slices of mice injected with AAV-GFAP-hPMCA2w/b-mCherry or AAV-GFAP-tdTomato control virus. A total of 542 events for control virus, and 292 events for hPMCA2w/b expression were analyzed (*n* = 2). Wilcoxon rank sum test. Comparison of signal amplitude, p = 0.0047; h = 1; zval: −2.8262, ranksum: 107058; duration (50%), p = 2.3276e-005; h = 1; zval: −4.2309, ranksum: 1.0304e+005; frequency: p = 21.2731e-033; h = 1; zval: −12.0847, ranksum: 7.7542e+004. **b**, Phenylephrine-evoked astrocyte Ca^2+^ signals in slices of mice injected with AAV-GFAP-hPMCA2w/b-mCherry or AAV-GFAP-tdTomato control virus. A total of 738 events for control virus, and 604 events for hPMCA2w/b expression were analyzed (*n* = 2). Wilcoxon rank sum test. Comparison of signal amplitude, p = 7.2e-034; h = 1; zval: −12.1319, ranksum: 319897; duration (50%), p = 5.8776e-015; h = 1; zval: −7.8065, ranksum: 350558. Data are means ± s.e.m.

**Extended Data Fig. 2.**
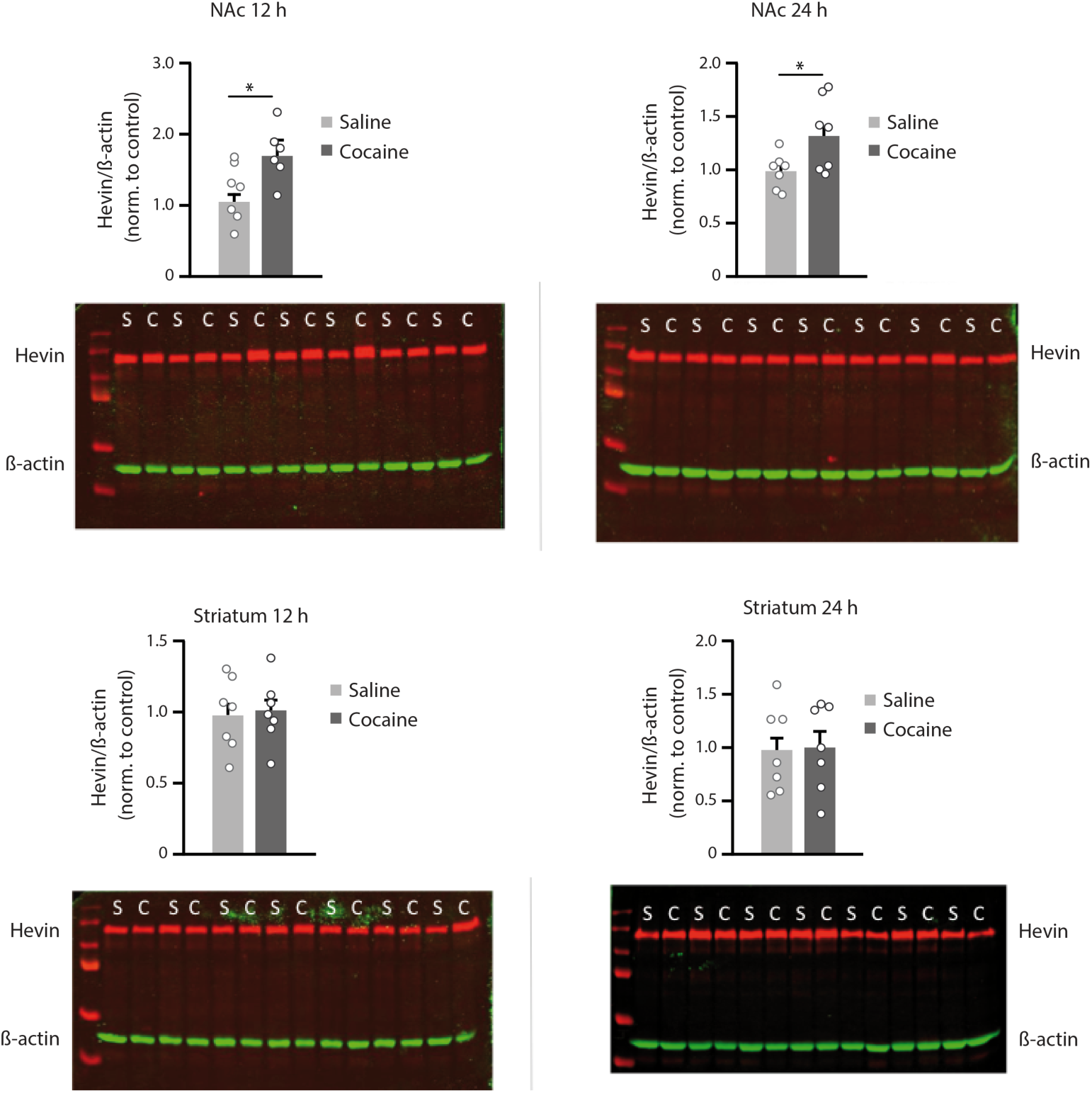
Effects of cocaine injection on hevin protein levels in the NAc and striatum. Western blot for hevin in NAc and dorsal striatum 12 and 24 h after a single cocaine injection (10 mg/kg, i.p.). Hevin expression was increased after 12 and 24 h only in NAc. No changes were observed in striatum. Hevin appears in green and ß-actin in red.

**Extended Data Fig. 3.**
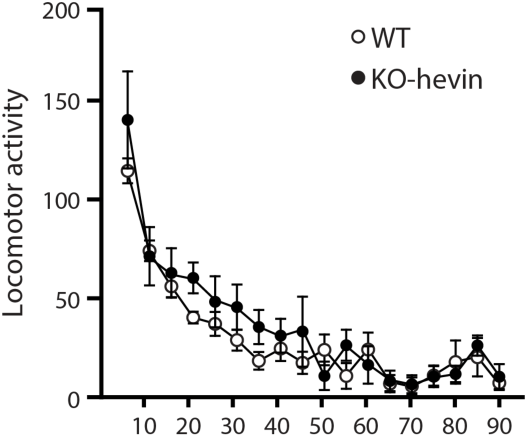
Locomotor activity was assessed in hevin-null mice and wild-type littermates in actimeter chambers, and divided in 5-min epochs over a period of 90 min.

**Extended Data Fig. 4.**
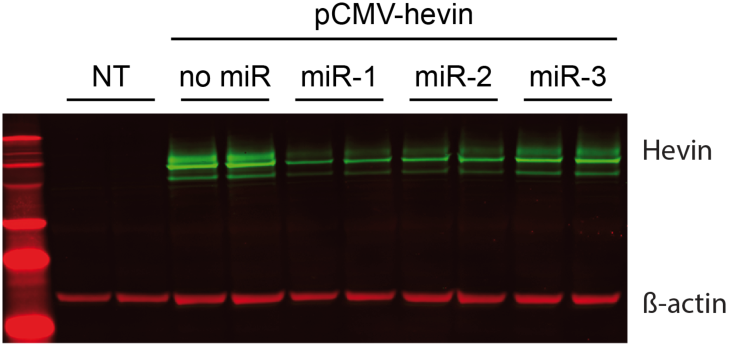
Hevin knockdown efficiency in Bon cells transfected with 3 distinct miR-hevin sequences. The three sequences were cloned in tandem into a single vector. Hevin appears in green and ß-actin in red. NT, non-transfected cells.

**Extended Data Fig. 5.**
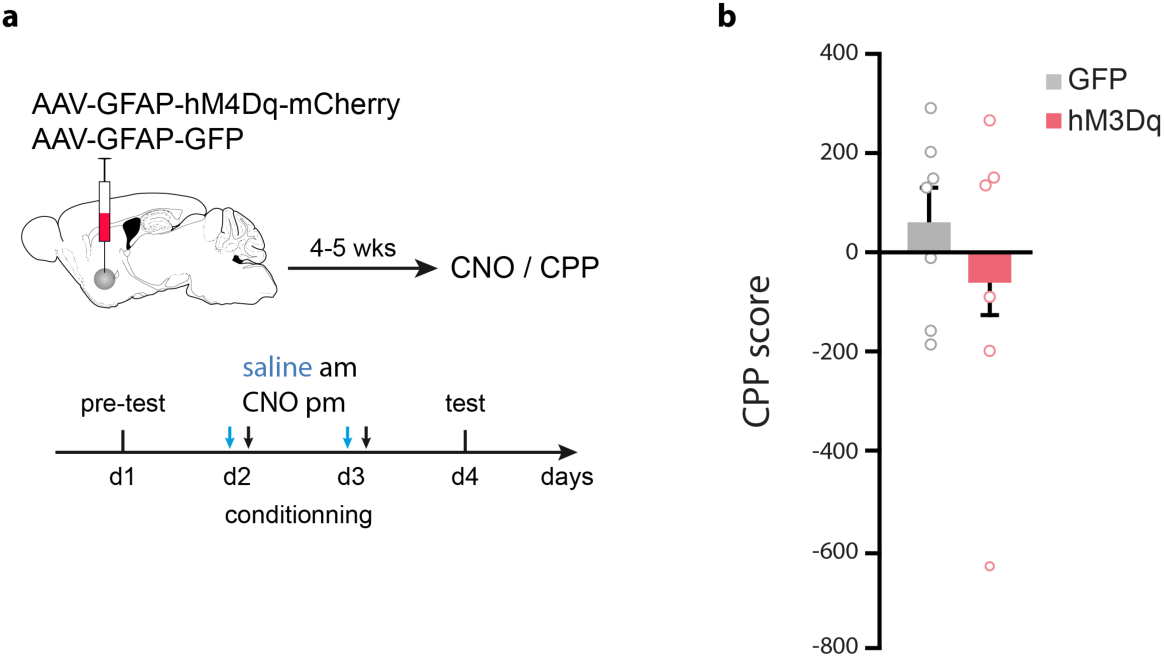
CPP to chemogenetic activation of astrocytes. **a**, Experimental procedure for chemogenetic activation of NAc astrocytes and timeline of CPP after CNO injection. **b**, CPP score after CNO injection paired with a compartment. CNO alone did not have any rewarding properties (*n* = 6-7). Data are means ± s.e.m.

**Extended Data Fig. 6.**
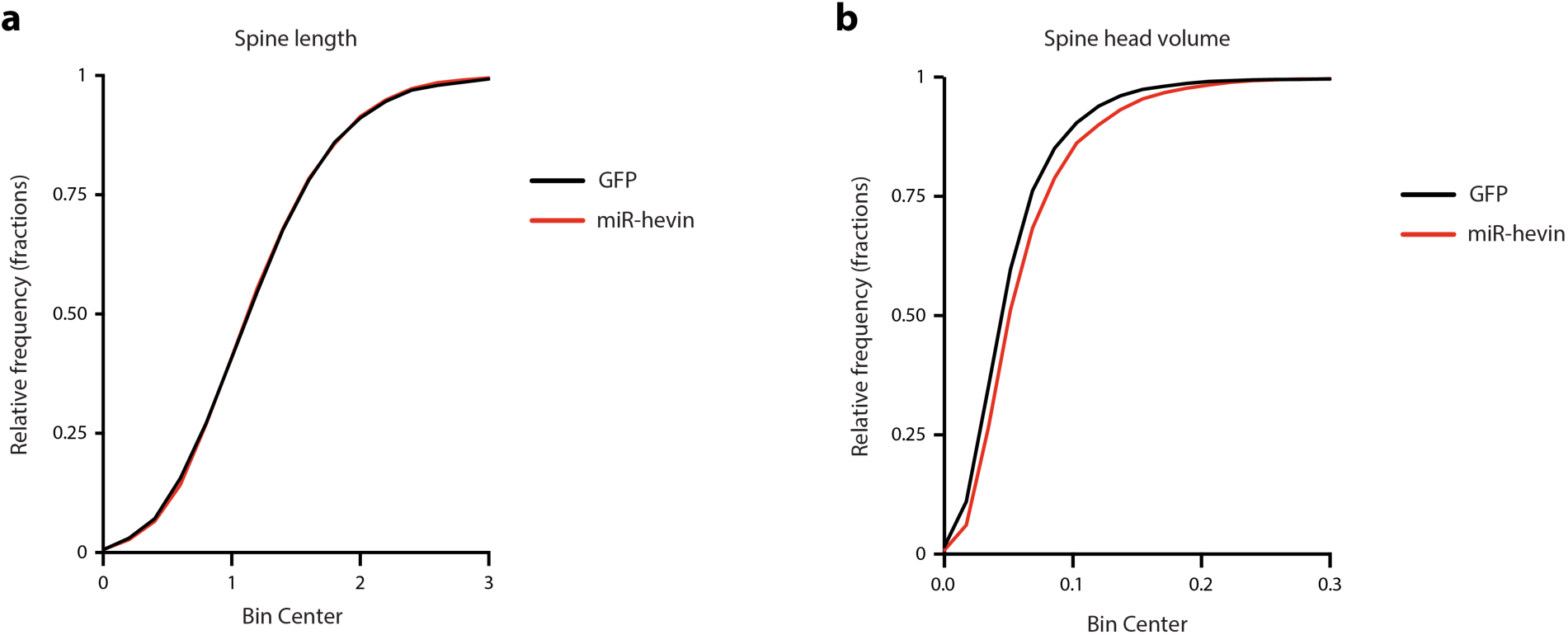
Dendritic spines in NAc MSN. **a** Spine length or **b**, spine head diameter of NAc MSN were not affected after hevin knockdown. *n* = 4-5.

**Extended Data Fig. 7.**
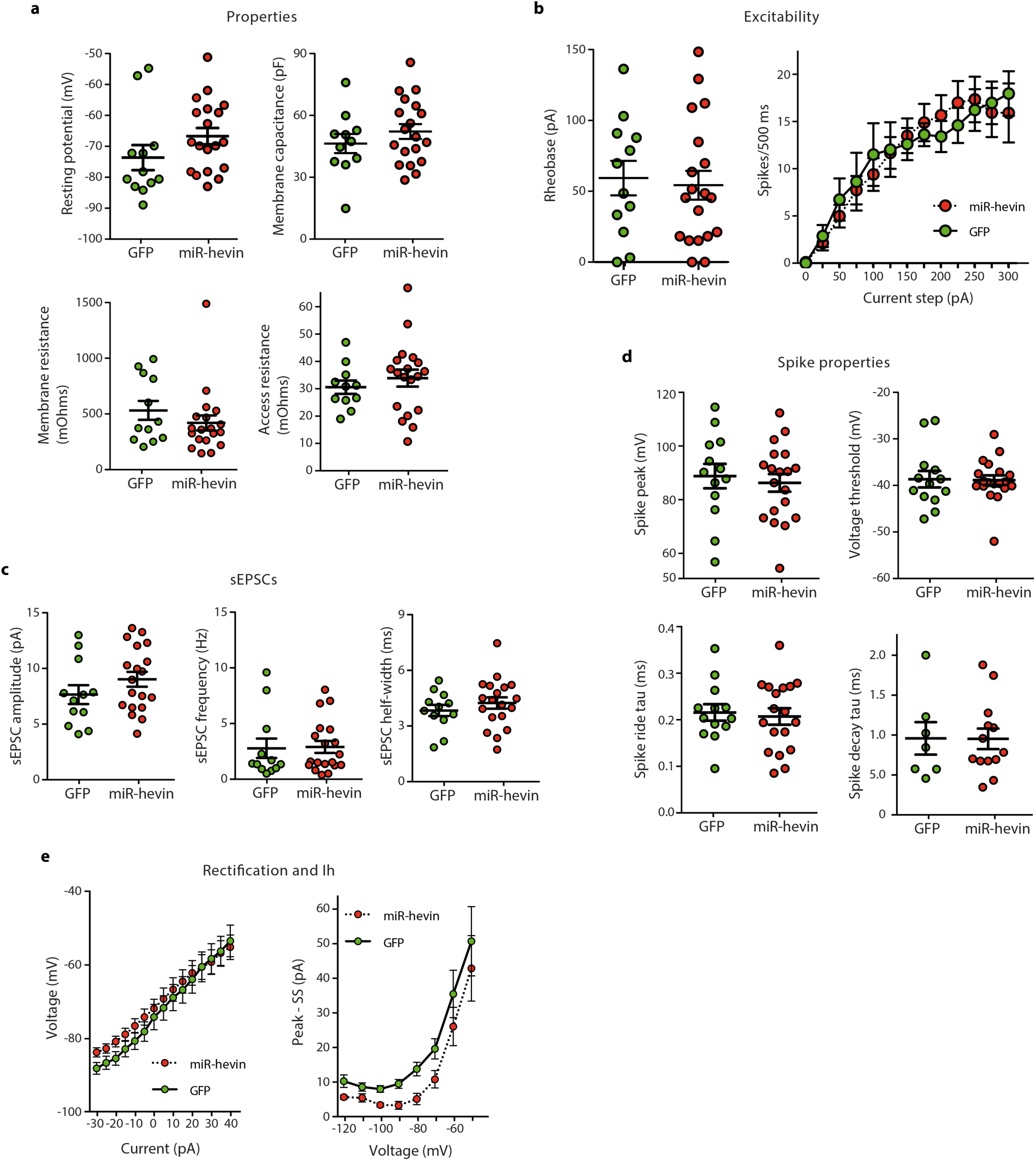
Intrinsic properties of NAc MSN at basal state were not affected by hevin knockdown in astrocytes. **a**, Resting membrane potential, membrane capacitance, membrane resistance, and access resistance in MSN of mice injected with AAV-GFAP-miR-hevin or AAV-GFAP-GFP control virus. **b**, Rheobase current and spike firing across depolarizing currents steps in MSNs. **c**, Spontaneous excitatory post-synaptic currents (EPSC) amplitude, frequency, and half-width in MSNs. **d**, Current-voltage plot and sag across voltage steps in MSNs. **e**, Spike properties in MSNs. Data are means ± s.e.m.

