## Supplementary material for "Astrocytes control cocaine-induced synaptic plasticity and reward through the matricellular protein hevin": Online Methods

### Animals

All animal experiments were carried out in compliance with the European Directive 2010/63/EU on the protection of animals used for scientific purposes, and approved by the local Animal Experimentation Ethics C2EA-05 Charles Darwin, the French Ministry of Higher Education and Research, and the Institutional Animal Care and Use Committee at Michigan State University in accordance with AAALAC. Seven to eight-week-old male C57BL/6J mice (Janvier Labs or Jackson Laboratory) were allowed at least 5 d to acclimate to the facility prior to any experimental procedures. All mice were housed in groups under standard conditions at 22±1°C and a 12 h light/dark cycle, with food and water provided *ad libitum*. Hevin-null (*Sparcl1*$^{-/-}$) mice were generated previously using homologous recombination of the second exon at the transcription start codon, resulting in the absence of hevin protein expression[1]. Hevin-null mice were backcrossed to C57BL/6J for 10 generations. Experiments were performed on 2 to 6-months-old hevin-null, wild-type (WT) littermates and C57BL/6J mice. For light-sheet imaging experiments, the genetically encoded $Ca^{2+}$ indicator GCaMP6f was expressed in astrocytes *in vivo* by crossing a Cre-dependent GCaMP6f mouse line (Ai95D, Jackson Laboratory) with an astrocyte-specific Glast-CreERT2 mouse line[2]. After generation of the bigenic GCaMP6f-GLAST mice, expression of Cre recombinase was induced by injecting tamoxifen (4 mg/kg, T5648, Sigma-Aldrich) daily for three consecutive days in 4 to 6-week-old mice[3].

### Drugs

Cocaine hydrochloride (Cooper Industries and Sigma-Aldrich) was dissolved in saline solution (0.9% NaCl). Clozapine N-oxide (CNO, R&D Systems, 4936-50) was dissolved in 25% DMSO at 10 mg/ml stock concentration, stored at -20°C, and protected from light. CNO was diluted daily in saline solution at a final concentration of 0.1 mg/ml. A ketamine and xylazine mixture (90 mg/kg and 10 mg/kg, respectively) diluted in saline solution was used as anesthesia for stereotaxic surgery. All drugs were administered intraperitoneally. Picrotoxin (Sigma-Aldrich, P1675) was dissolved in 100% DMSO and diluted to 0.1% DMSO in artificial cerebrospinal fluid (aCSF). D-AP5 (HelloBio, HB0225) was dissolved in double-distilled $H_2O$.

### Cocaine conditioned place preference (CPP)

CPP was performed using two distinct chambers connected by a middle neutral chamber (Imetronic) as described previously[4]. The two chambers have distinct visual and tactile cues to allow differentiation. During the pre-test (preconditioning phase), mice were allowed to freely explore both chambers for 30 min. Assignation of the cocaine-paired chamber was adjusted to balance out any pre-existing chamber bias. In the conditioning phase, mice were injected with saline (morning) or cocaine (afternoon) and placed for 20 min in a given chamber for two consecutive days. For the experiments using chemogenetic activation of astrocyte $Ca^{2+}$ signals, hM3Dq- and GFP-mice were injected with CNO (1 mg/kg) in their home cage 30 min prior to the start of conditioning to cocaine. On the test day, animals were allowed to explore both chambers for 30 min. The time spent in each chamber during the test and pre-test was recorded. The place preference score was determined as the time spent on the cocaine-paired chamber during the test minus the pre-test. The rewarding properties of CNO alone were tested by injecting CNO 30 min prior to placing the mice into the CNO-paired chamber without cocaine injection.

**Locomotor activity**

Locomotor activity was measured using a circular corridor with four infrared beams placed at every 90° (Imetronic) in a low-luminosity environment. Locomotor activity was counted as travels through one-fourth of the circular corridor as detected by consecutive interruption of two adjacent beams. Mice were placed in the activity apparatus for 90 min.

**Design of microRNA directed against hevin mRNA**

Synthetic microRNAs (miR) against hevin were designed, validated *in vitro* and cloned into the adeno-associated virus AAV2/5-GFAP-eGFP. More precisely, oligonucleotides targeting hevin mRNA were selected using Dharmacon online resources (Dharmacon, ThermoFisher). Three sequences were selected based on selectivity and tested in BON cells co-transfected with hevin-eGFP expressing vector. Based on hevin protein knockdown efficiency, three shRNA were chosen for miR engineering (BLOCK-iT Pol II miR RNAi Expression Vector Kits, Invitrogen, K4935-00) following the vendor's instructions. Top target pre-miRNA and complementary bottom strand oligonucleotides were designed and annealed to generate double-stranded oligonucleotides and cloned into pcDNA 6.2-GW/EmGFP. RNA interference efficacy was tested using BON cells transfected with full-length hevin inserted into pCMV-SPORT6 vector. Three miRNA sequences following the EmGFP sequence were PCR-amplified and cloned in tandem (HiFi assembly, New England Biolabs, E2621) into the pAAV2-GFAP-eGFP viral vector (Addgene, 50473) by replacing the eGFP sequence. The human GFAP promoter (GfaABC1D) is highly selective for astrocytes[5]. A negative scramble miRNA control sequence was cloned into pAAV2-GFAP-eGFP. Viral preparations were made at Atlantic Gene Therapy and the AAV production facility at Institut de la Vision. The knockdown efficiency of hevin was validated *in vivo*.

**Stereotaxic surgery**

Eight-week-old mice were anesthetized using a mix of ketamine and xylazine (90 mg/kg and 10 mg/kg diluted in saline), and placed in a stereotaxic frame (Kopf Instruments). Lidocaine was applied at the location of the scalp incision before the procedure. Meloxicam (Metacam, 1 mg/kg subcutaneously) was injected before and after surgery for peri- and post-operative pain relief. Burr holes were then drilled in the skull in accordance with the planned injection coordinates (see below). All mice were allowed to recover from the surgery for at least 7 days before the experiments.

**Virus injections**

Viral vectors were injected using a 5 µl microsyringe (Hamilton) at a rate of 0.1 µl/min. AAV2/5-GFAP-EmGFP-miR-hevin ($9.4 \times 10^{12}$ vg/ml, Atlantic Gene Therapy, Université de Nantes, INSERM), AAV2/5-GFAP-HA-hM3D(Gq)-IRES-mCitrine ($4 \times 10^{12}$ vg/ml, University of North Carolina), AAV2/5-GFAP-mCherry-hPMCA2w/b (Addgene, 111568, viral preparation: $7 \times 10^{13}$ vg/ml, Institut de la Vision) and control viruses AAV2/5-GFAP-eGFP-WPRE-hGH ($9.1 \times 10^{12}$ vg/ml, University of Pennsylvania viral core) and AAV2/5-GFAP-EmGFP-NegativeControl-miRNA ($1 \times 10^{14}$ vg/ml, Institut de la Vision) were injected bilaterally into the nucleus accumbens (NAc) of adult C57BL/6J mice with the following coordinates: +1.5 AP, +1.45 ML, -4.3 DV mm, relative to bregma, 10° angle. For $Ca^{2+}$ fiber photometry experiments, mice were injected unilaterally with AAV2/5-GFAP.cyto-GCaMP6f ($1.4 \times 10^{13}$ GC/ml, Addgene, 52925) into the NAc. For chemogenetics astrocyte activation and $Ca^{2+}$ signals recording, a mixture of AAV2/5-GFAP-hM3D(Gq)-mCherry ($1.3 \times 10^{13}$ GC/ml, Addgene, 50478) with AAV2/5-GFAP.cyto-GCaMP6f was injected. Behavioral analyses and imaging experiments were performed 4-6 weeks after the surgery. Injection sites and viral transgene expression were confirmed for all mice by examination of endogenous eGFP fluorescence, or after immunohistochemistry for mCherry, GCaMP6f or mCitrine.

**Immunohistochemistry**

Mice were anesthetized and perfused intracardially with 4% paraformaldehyde (PFA) in phosphate buffer solution (PBS). Brains were removed and left at 4°C in 4% PFA overnight, then kept in PBS with azide (0.01%) until use. Brain slices (30 µm) were then cut on a vibratome and kept in PBS with azide at 4ºC until use. Slices were incubated in a blocking buffer containing 0.2% bovine serum albumin and 0.5% triton for 1 h at room temperature. Slices were then treated with antibodies against hevin (1/1500, R&D, mSPARCL1 AF2836), NeuN (1/1000, Aves Labs, NUN-9889), GFAP (1/1000, Chemicon, MAB360), GFP (1/1000, Invitrogen, A11122), or RFP (1/1000, Rockland, 600-401-379) overnight at 4°C. After washing, slices were incubated with a CY2 or CY3 (1/4000, Jackson) conjugated secondary antibody for 2 h at room temperature, and then washed with PBS 3 times for 15 min each. Nuclear staining was performed with 4' 6-diamidino-2-phenylindole (DAPI). Sections were mounted on slides and coverslipped with Fluoromount (Sigma-Aldrich, F4680) for fluorescent image acquisition.

**Image acquisition**

All slides were scanned on a NanoZoomer 2.0-HT (Hamamatsu Photonics) at 20x resolution. Laser intensity and time of acquisition was set separately for each antibody. Images were analyzed using the NDP.view2 software (Hamamatsu Photonics). Regions of interest were identified according to the Paxinos mouse brain atlas[6]. Positive cells refer to a staining in a cell body clearly above background and surrounding a DAPI-stained nucleus. Co-localization was determined by the presence of the signals for both probes in the soma of the same cell. The entire region of interest was evaluated. For illustration purposes, the NanoZoomer images were exported in TIFF format using NDP viewer. Images were corrected for contrast, cropped on Photoshop CS6, and assembled on Illustrator CS6 (Adobe).

**Western blot**

Whole tissue extracts were prepared from bilateral punches (1–1.5 mm diameter; Miltex) of brain regions from 10 week-old mice treated with cocaine. Samples were homogenized by sonication on ice in 200 µl of RIPA buffer containing a protease inhibitor cocktail (1/1000, Sigma-Aldrich, 11836153001). Protein concentrations were determined by Bradford's method and the samples were stored at -70ºC until analyzed by Western blot. Protein samples (50 µg) were prepared with NuPage LDS sample buffer (Invitrogen, NP0008) and DTT 1 M 10%, and heated at 90°C for 10 min. Samples were loaded and separated by Bis-Tris sodium dodecyl sulfate polyacrylamide gel electrophoresis (10% gels) and transferred onto PVDF membranes (Merck, Immobilon IPFL85R). Transfer efficacy was controlled by Ponceau staining. Unspecific binding sites were blocked with Tris-buffered saline containing 5% nonfat milk and 0.1% Tween-20, and membranes were immunoprobed with antibodies against hevin (1/1000, R&D, mSPARCL1 AF2836) and ß-Actin (1/5000, Sigma, A1978). Membranes were incubated with infrared-labeled secondary antibodies (IRDye 680DX and IRDye 800CW; 1/5000; LI-COR Biosciences) in phosphate buffer containing 5% nonfat milk. Immunoblotting was quantified with the Odyssey Infrared Imaging System and Application Software version 3.0 (LI-COR Biosciences).

**Diolistic labeling of neurons**

Dendritic spines were labeled in brain sections from perfused mice using the Diolistic technique. Briefly, 50 mg of tungsten beads were mixed with 3 mg of solid red DiI (3,3'-dioctadecyloxacarbocyanine perchlorate) dissolved in methylene chloride. Cartridges were prepared by coating the beads on the inner surface of a Teflon tube pretreated with polyvinylpyrrolidone. Helium gas pressure was applied with a gene-gun device to project the beads out of the cartridge onto the brain section. The beads were delivered through a 3-micron pore-size filter to avoid beads clusters. After labeling, the sections were kept in PBS at room temperature for at least 2 h and mounted in Prolong Gold (ThermoFisher, P36930).

**Confocal microscopy and analysis of spine density**

Image stacks were taken using a confocal laser scanning microscope (SP5, Leica) equipped with a 1.4 NA objective (oil immersion, Leica) with a pinhole aperture set to 1 Airy unit, pixel size of 60 nm and z-step of 200 nm. The excitation wavelength and emission range were 488 and 500–550 for GFP, 561 and 570–620 for DiI. Metrology measurements were regularly performed using fluorescent beads to confirm proper laser alignment, laser power and field homogeneity using the ImageJ-based MetroloJ plugin. Deconvolution with experimental point spread function from fluorescent beads using a maximum likelihood estimation algorithm was performed with Huygens software (Scientific Volume Imaging). NeuronStudio software was used to reconstruct the dendrite and detect dendritic spines in 3D. When necessary, manual correction was applied. Dendritic spine density was defined as the number of spines normalized to a 10 µm length of dendrite. Spine heads were segmented in 3D with a custom procedure described in [7].

**Electrophysiological recordings**

Stereotaxic surgery was conducted as previously described. Viral vectors (AAV2/5-GFAP-EmGFP-miR-hevin; AAV2/5-GFAP-eGFP) were bilaterally infused into the NAc. Mice were maintained in their home cage throughout the duration of cocaine administration. Cocaine (10 mg/kg; i.p.) or saline was injected once daily for 5 d. *Ex vivo* electrophysiology experiments were conducted 1 d after the last cocaine injection. Brains from transcardially perfused mice were sliced into coronal sections (250 µM containing NAc) in sucrose aCSF (in mM: 234 sucrose, 2.5 KCl, 1.25 $NaH_2PO_4$, 10 $MgSO_4$, 0.5 CaCl, 26 $NaHCO_3$, and 11 glucose; bubbled with 95% $O_2$-5% $CO_2$). Slices were then transferred to an incubation chamber containing aCSF (in mM: 126 NaCl, 2.5 KCl, 1.25 $NaH_2PO_4$, 2 MgCl, 2 CaCl, 26 $NaHCO_3$, and 10 glucose; bubbled with 95% $O_2$-5% $CO_2$) and held at 34°C. Recordings were made from a submersion chamber perfused with aCSF (2 ml/min) maintained at 32-34°C. Borosilicate glass electrodes (3–6 MΩ) were filled with Cs-methanosulfate internal solution (in mM: 110 cesium methanosulfate, 20 TEA Cl, 8 KCl, 10 HEPES, 0.2 EGTA, 2 Mg-ATP, and 0.4 Na-GTP; pH 7.2–7.4, 290 mOsm). NAc medium spiny neurons (MSNs) in the milieu of GFP-positive glia were visualized using an upright DIC microscope (Olympus) using infrared and epifluorescent illumination. NAc MSNs were distinguished by location and individual cell morphology. Whole-cell patch-clamp recordings were made from NAc MSNs using a Multiclamp 700B amplifier and a Digidata 1440A digitizer (Molecular Devices). Whole-cell junction potential was not corrected. Traces were sampled (20 kHz), filtered (Bessel 2 kHz), and digitally stored. Cells with series resistance greater than 30 MΩ were omitted from analysis. For evoked EPSC recordings, a monopolar stimulating electrode was placed 200-500 µm away from the patched neuron and another stimulating electrode was placed into the bath. Stimulation intensity varied from 100-2000 µA for 10 µs. EPSCs (15-30 traces; 10 s inter-stimulus interval) were recorded while the cell was held at -70 mV (for AMPA-R EPSCs) and +40 mV (for NMDA-R EPSCs). For paired pulse recordings, paired EPSCs (10 traces) were evoked at -70 mV with inter-stimulus intervals of 20 ms, 50-500 ms, and 1 s. Traces were compiled in pClamp 10 (Clampfit). Baseline was adjusted to the first 100 ms prior to stimulation. For assessment of AMPA-R EPSC at -70 mV, the averaged peak negative-going current within the first 15 ms was recorded. For assessment of NMDA-R EPSCs at +40 mV, the averaged current at 25 ms post peak was recorded.

**Fiber photometry experiments**

Eight-week-old WT mice were injected unilaterally with 0.5 µl of AAV2/5-GFAP.cyto-GCaMP6f (1.4x10e13 GC/ml, Addgene) in the NAc (+1.5 AP, ±1.45 ML, -4.3 DV mm relative to bregma, 10° angle). Optical fibers (200 mm core, NA = 0.37, Neurophotometrics) coupled to a stainless-steel ferule (1.25 mm) were implanted after virus injection at the same site (coordinates: +1.5 AP, ±0.8 ML, -4.3 DV mm relative to bregma, without angle), and fixed to the skull with dental cement (SuperBond, Sun Medical). Two weeks after surgery, animals began a habituation period to the open-field used for photometry recordings and handling. All recordings were performed in a novel cage between 10:00 and 16:00. Mice were habituated to the cage and fiber setup for 30 min, and the $Ca^{2+}$ activity was recorded for 55 min (15 min of baseline, 15 min after saline injection and 25 min after cocaine injection). Fluorescence measurements of $Ca^{2+}$ signals were recorded using a fiber photometry system (Neurophotometrics, FP3002). A branched fiber optic patch cord (BFP_200/230/900-0.37_FC-MF1.25, Doric Lenses) connected to the fiber photometry apparatus was attached to the implanted fiber optic cannula using a cubic zirconia sleeve. To record fluorescence signals from GCaMP6f, light from a 470 nm LED was band-pass filtered, collimated, reflected by a dichroic mirror and focused by a ×20 objective. LED light was delivered at a power that resulted in 50 µW of 470 nm light at the tip of the patch cord. Emitted GCaMP6f fluorescence was band-pass filtered and focused on the sensor of a CCD camera. To account for auto-fluorescence and possible motion artifacts during testing, a 410 nm light stimulation corresponding to isosbestic GCaMP6f signal was used. This signal was similarly directed into the brain and subsequently measured with the CCD camera. Signals were collected at a rate of 40 Hz and visualized using the open-source software Bonsai 2.4 (http://bonsai-rx.org).

**Fiber photometry analysis**

Data analyses were performed using a Matlab script[8]. GCaMP6f signals recorded at 410 nm are not $Ca^{2+}$ dependent; thus, change in signal can be attributed to autofluorescence, bleaching and fiber bending. Accordingly, to calculate ΔF/F, a linear fit was applied to the 410 nm signal to align it to the 470 nm signal, producing a fitted 410 nm. This signal was used as F0 to normalize the signal at 470 nm using standard ΔF/F normalization ([470 nm signal - fitted 410 nm signal]/fitted 410 nm signal). For the chemogenetic experiments, the average signal in the 5 min preceding the i.p. injection (baseline) was compared to the average signal 5 min after the injection. For the cocaine experiments, the peak oscillation pattern was analyzed using the GuPPy software[9]. All data were downsampled to 4 Hz. The first and last minute of recording were removed, along with artifacts, so the remaining trace was related to the $Ca^{2+}$ activity, and not to signal fluctuation. Peak detection used a 15, 20 and 25 s moving window. Detected peaks were validated visually by a blinded experimenter. High amplitude events (events with amplitudes greater than two times the median absolute deviation - MAD - above the median of the moving window) are filtered out and the trace is recalculated; local maxima higher than three MADs are considered a signal. We compared the signal 5 min before i.p. injection (baseline) to the signal between 5 and 15 min after i.p. injection (saline or cocaine). The injection site and fiber tip localization were analyzed for all mice after the experiments.

**Light-sheet imaging**

Light-sheet fluorescent microscopy enables wide field-optical sectioning, thereby high-quality dynamic imaging of a thin layer of the slices. Light sheet imaging was performed using a custom-built setup based on a wide-field upright microscope (Zeiss Axioskop 50) equipped with water-immersion objectives. The in-focus light sheet was generated from an independent optical module equipped with an air objective (10X 0.25NA, Zeiss) and orthogonally positioned to the upright imaging pathway. This module was coupled via a wavelength combiner (Thorlabs) to two continuous wave lasers (473 nm and 561 nm, CNI). GCaMP6f was excited by the 473 nm laser via a double-band dichroic (Di03-R488/561-t3, Semrock) and fluorescence was collected via the emission filter (FF01-523/610, Semrock) and a digital electron-multiplying charged-coupled device (EMCCD Cascade 512B, Photometrics). All devices were controlled by MetaMorph Software (Molecular Devices). During imaging, the center of the light sheet was orthogonally aligned to the focus of the imaging objective. To allow free access to any parts of the brain slice, an angular light sheet system was developed. Briefly, before imaging, the brain slice was carefully laid on a custom-designed chamber and stabilized with a harp slice anchor grid (Warner Instruments). The horizontal surface supporting the slice had an inclination of 8° along the laser entry direction. The chamber was mounted on a motorized stage (Physik Instrumente) for axial micro-manipulation. Time-lapse images were acquired at 1 Hz with 300 ms exposure time. The effect of hPMCA2w/b on astrocytic $Ca^{2+}$ signal dynamics was validated using light-sheet microscopy and analyzed by the event-based method AQuA[10]. An AAV2/5-GFAP-mCherry-hPMCA2w/b was injected unilaterally into the NAc of GCaMP6f-GLAST double transgenic mice. The 300 µm-thick coronal slices comprising NAc were acutely prepared from GCaMP6f-GLAST mice of both sexes. The brains were quickly taken out and placed in an ice-cold modified aCSF (maCSF, in mM: 30 NaCl, 4.5 KCl, 1.2 NaH2PO4, 1 MgCl2, 26 HaHCO3, 10 D-Glucose and 194 sucrose) and cut into slices with a vibratome (Leica VT1200S) in oxygenated mACSF maintained at 4°C. Brain slices were left to recover 1 h in normal aCSF (in mM: 124 NaCl, 4.5 KCl, 1.2 NaH2PO4, 1 MgCl2, 2 CaCl2, 26 NaHCO3, 10 D-Glucose) at 37°C with constant oxygenation. Imaging was carried out at room temperature (21–23°C) under constant perfusion (~3 ml/min) with normal aCSF. Solutions were saturated with a mixture of 95% O2 and 5% CO2 for all steps.

**Statistics**

PRISM (GraphPad software) was used for statistical calculations. The data generated during this study is detailed in Table S1. Results were first tested for normality using D'Agostino-Pearson omnibus normality test, and F-test for equality of variances. Student's t-test was used when parameters followed a normal distribution, otherwise data were analyzed with a non-parametric Mann-Whitney test. One-way or two-way repeated ANOVA followed by Tukey's post hoc comparison was used for normally distributed data. Animals were excluded from experimental data only when they were identified as outliers by the outlier test (Grubb's test, GraphPad software). All data are expressed as mean ± standard error of the mean (s.e.m.), and statistical differences were established with $p < 0.05$, $p < 0.01$ and $p < 0.001$. For electrophysiological experiments, the AMPA/NMDA ratio was assessed by the difference (relative to saline GFP control mice) between AMPA-R EPSCs and NMDA-R EPSCs. Paired pulse ratio (PPR) was assessed by the difference between the 2$^{nd}$ EPSC compared to the 1$^{st}$ EPSC in the mean peak negative-going current within 15 ms following stimulation. Electrophysiological data were analyzed by one-way or two-way ANOVAs followed by Holm-Sidak corrected post hoc comparisons.
